# Decoding sequence-level information to predict membrane protein expression

**DOI:** 10.1101/098673

**Authors:** Shyam M. Saladi, Nauman Javed, Axel Müller, William M. Clemons

## Abstract

The expression of integral membrane proteins (IMPs) remains a major bottleneck in the characterization of this important protein class. IMP expression levels are currently unpredictable, which renders the pursuit of IMPs for structural and biophysical characterization challenging and inefficient. Experimental evidence demonstrates that changes within the nucleotide or amino-acid sequence for a given IMP can dramatically affect expression; yet these observations have not resulted in generalizable approaches to improved expression. Here, we develop a data-driven statistical predictor named IMProve, that, using only sequence information, increases the likelihood of selecting an IMP that expresses in *E. coli.* The IMProve model, trained on experimental data, combines a set of sequence-derived features resulting in an IMProve score, where higher values have a higher probability of success. The model is rigorously validated against a variety of independent datasets that contain a wide range of experimental outcomes from various IMP expression trials. The results demonstrate that use of the model can more than double the number of successfully expressed targets at any experimental scale. IMProve can immediately be used to identify favorable targets for characterization.

## Introduction

The biological importance of integral membrane proteins (IMPs) motivates structural and biophysical studies that require large amounts of purified protein at considerable cost. Only a small percentage can be produced at high-levels resulting in IMP structural characterization lagging far behind that of soluble proteins; IMPs currently constitute less than 2% of deposited atomic-level structures ^1^. To increase the pace of structure determination, the scientific community created large government-funded structural genomics consortia facilities, like the NIH-funded New York Consortium on Membrane Protein Structure (NYCOMPS)^2^. For this representative example, more than 8000 genes, chosen based on characteristics hypothetically related to success, yielded only 600 (7.1%) highly expressing proteins ^3^ resulting to date in 34 (5.6% of expressed proteins) unique structures (based on annotation in the RCSB PDB ^4^). This example highlights the funnel problem of structural biology, where each stage of the structure pipeline eliminates a large percentage of targets compounding into an overall low rate of success ^5^. With new and rapidly advancing technologies like cryo-electron microscopy, serial femtosecond crystallography, and micro-electron diffraction, we expect that the latter half of the funnel, structure determination, will increase in success rate ^6–8^. However, IMP expression will continue to limit targets accessible for study ^9^.

Tools for improving the number of expressed IMPs are needed. While significant work has shown promise on a case-by-case basis, *e.g.* growth at lower temperatures, codon optimization ^10^, and regulating transcription ^11^, a generalizable solution remains elusive. Currently, each target must be addressed individually as the conditions that were successful for a previous target seldom carry over to other proteins, even amongst closely related homologs ^5,2^. For individual cases, simple changes can have dramatic effects on the amount of expressed proteins ^13,14^ Considering the scientific value of IMP studies, it is surprising that there are no methods that can provide solutions for improved expression outcomes with broad applicability across protein families and genomes.

There are currently no approaches available that can decode sequence-level information for predicting IMP expression; yet it is common knowledge that sequence changes which alter overall biophysical features of the protein and mRNA transcript can measurably influence IMP biogenesis. While physics-based approaches which have proven successful in correlating integration efficiency and expression ^12,15^, that and other work revealed that simple application of specific ‘sequence features’, such as the positive-inside rule, are inadequate to predict IMP expression ^16,17^. For the positive-inside rule, as an example, this contrasts evidence that the number of positive-charges on cytoplasmic loops is known to be an important determinant of IMP biogenesis ^18,19^. The reasons for this failure to connect sequence to expression likely lie in the complex underpinnings of IMP biogenesis, where the interplay between many sequence features at both the protein and nucleotide levels must be considered. Optimizing for a single sequence feature likely diminishes the beneficial effect of other features (*e.g.* increasing positive residues on internal loops might diminish favorable mRNA properties). Without accounting for the broad set of sequence features related to IMP expression, it is impossible to predict differences in expression.

Development of a low-cost, computational resource that significantly and reliably predicts improved expression outcomes would transform the study of IMPs. Attempts to develop such algorithms have so far failed. Several examples, Daley, von Heijne, and coworkers ^10,16,17^ as well as NYCOMPS, were unable to use experimental expression data sets to train models that returned any predictive performance (personal communication). This is not surprising, given the difficulty of expressing IMPs and the limits in the knowledge of the sequence features that drive expression. In other contexts, statistical tools based on sequence have been shown to work; for example, those developed to predict soluble protein expression and/or crystallization propensities ^20–22^. Such predictors are primarily based on available experimental results from the Protein Structure Initiative ^23,24^. While collectively these methods have supported significant advances in biochemistry, none of the models are able to predict IMP outcomes due to limitations inherent in the model development process. As IMPs have an extremely low success rate, they are either explicitly excluded from the training process or are implicitly down-weighted by the statistical model (for representative methodology see ^25^). Consequently, none have successfully been able to map IMP expression to sequence.

Here, we demonstrate for the first time that it is possible to predict IMP expression directly from sequence. The resulting predictor allows one to enrich expression trials for proteins with a higher probability of success. To connect sequence to prediction, we develop a statistical model that maps a set of sequences to experimental expression levels via calculated features—thereby simultaneously accounting for the many potential determinants of expression. The resulting IMProve model allows ranking of any arbitrary set of IMP sequences in order of their relative likelihood of successful expression. The IMProve model is extensively validated against a variety of independent datasets demonstrating that it can be used broadly to predict the likelihood of expression in *E. coli* of any IMP. With IMProve, we have built a way for more than two-fold enrichment of positive expression outcomes relative to the rate attained from the current method of randomly selecting targets. We highlight how the model informs on the biological underpinnings that drive likely expression. Finally, we provide direct examples where the model can be used for a typical researcher. Our novel approach and the resulting IMProve model provide an exciting paradigm for connecting sequence space to complex experimental outcomes.

## Results

For this study, we focus on heterologous expression in *E. coli*, due to its ubiquitous use as a tool for expression across the spectrum of the membrane proteome. For example, 43 of the 216 unique eukaryotic IMP structures were solved using protein expressed in *E. coli* (based on annotation in the RCSB PDB ^4^). Low cost and low barriers for adoption highlight the utility of *E. coli* as a broad tool if the expression problem can be overcome.

### Development of a computational model trained on *E. coli* expression data

A key component of any data-driven statistical model is the choice of dataset used for training. Having searched the literature, we identified two publications that contained quantitative datasets on the IPTG-induced overexpression of *E. coli* polytopic IMPs in *E. coli.* The first set, Daley, Rapp *et al.,* contained activity measures, proxies for expression level, from C-terminal tags of either GFP or PhoA (alkaline phosphatase)^16^. The second set, Fluman *et al.,* used a subset of constructs from the first and contained a more detailed analysis utilizing in-gel fluorescence to measure folded protein ^26^ (see Methods 4c). The expression results strongly correlated (Spearman’s *ρ* = 0.73) between the two datasets demonstrating that normalized GFP activity was a good measure of the amount of folded IMP (Fig. 1A and ^26,27^). The experimental set-up employed multiple 96-well plates over multiple days resulting in pronounced variability in the absolute expression level of a given protein between trials. Daley, Rapp *et al*. calculated average expression levels by dividing the raw expression level of each protein by that of a control protein on the corresponding plate.

**Fig. 1.**
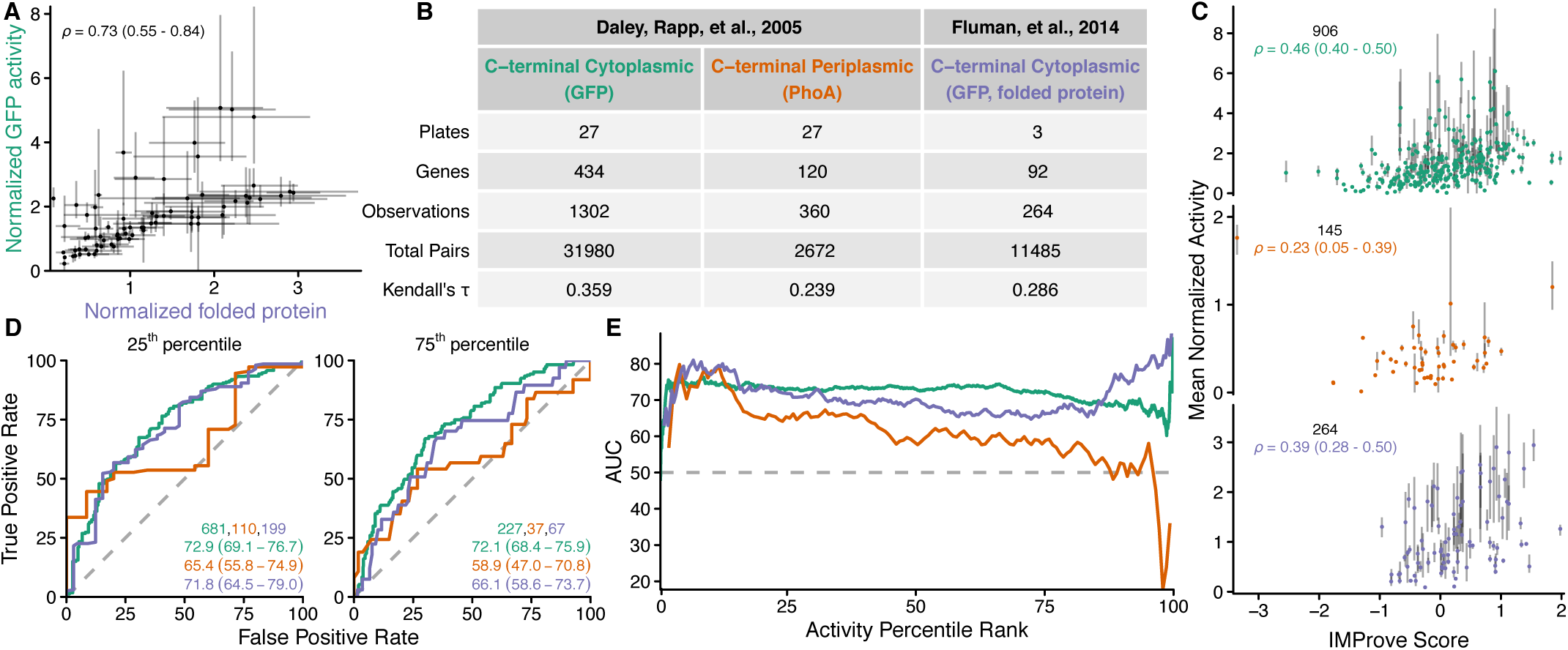
Training performance. **(A)** A comparison of GFP activity ^16^ with measured folded protein ^26^ where each point represents the mean for a given gene tested in both works, and error bars plot the extrema. Spearman’s rank correlation coefficient and 95% confidence interval (CI) ^104^ are shown. **(B)** Plates are the number of independent sets of measurements within which expression levels can be reliably compared. Genes are the number of proteins for which the C-terminus was reliably ascertained ^16^. Observations are the total number of expression data points accessible. Total pairs are the number of comparable expression measurements *(i.e.* those within a single plate). Kendall’s *t* is the metric maximized by the training process (See Methods 4b). The color of the column heading identifying each experimental set is retained throughout the figure. **(C)** Agreement against the normalized outcomes plotted as the mean activity (see Methods 5 for definition) versus the score with error bars providing the extent of observed activities (Spearman’s p and 95% CI noted). **(D)** Illustrative Receiver Operating Characteristics (ROC) for thresholds at 25^th^ and 75^th^ percentile in activity with the number of positive outcomes at that threshold, the Area Under the Curve (AUC), and 95% CI indicated. **(E)** The AUC of the ROC at every possible activity threshold.

To successfully map sequence to expression, we additionally needed to derive numerical features from a given gene sequence that are empirically related to expression. Approximately 105 sequence features from protein and nucleotide sequence were calculated for each gene using custom code together with published software (codonW ^28^, tAI ^29^, NUPACK ^30^, Vienna RNA ^31^, Codon Pair Bias ^32^, Disembl ^33^, and RONN ^34^). Relative metrics (e.g. codon adaptation index) are calculated with respect to the *E. coli* K-12 substr. MG1655^35^ quantity. The octanol-water partitioning ^36^, GES hydrophobicity ^37^, ΔG of insertion ^38^ scales were employed as well. Transmembrane segment topology was predicted using Phobius constrained for the training data and Phobius for all other datasets ^39^. Two RNA secondary structure metrics were prompted in part by Goodman, et al. ^40^. Supplementary Table 1 includes a detailed description of each feature. All features are calculated solely from the coding region of each gene of interest excluding other portions of the open reading frame and plasmid *(e.g.* linkers and tags, 5′ untranslated region, copy number).

Fitting the data to a simple linear regression provides a facile method for deriving a weight for each feature. However, using the set of sequence features, we were unable to successfully fit a linear regression using the normalized GFP and PhoA measurements reported in the Daley, Rapp *et al.* study. Similarly, using the same feature set and data, we were unable to train a standard linear Support Vector Machine (SVM) to predict the expression data either averaged or across all plates (see Supplementary Table 1; Methods 2,3). Due to the attempts by others to fit this data, this outcome may not be surprising and suggested that a more complex analysis was required.

We hypothesized that training on relative measurements across the entire dataset introduced errors that were limiting. To address this, we instead only compare measurements within an individual plate, where differences between trials are less likely to introduce errors. To account for this, a preference-ranking linear SVM algorithm (SVM^rank 41^) was chosen (see Methods 4b). Simply put, the SVM^rank^ algorithm determines the optimal weight for each sequence feature to best rank the order of expression outcomes within each plate over all plates, which results in a model where higher expressing proteins have higher scores. The outcome is identical in structure to a multiple linear regression, but instead of minimizing the sum of squared residuals, the SVM cost function accounts for the plate-wise constraint specified above. In practice, the process optimizes the correlation coefficient Kendall’s *τ* (Eq. 1) to converge upon a set of weights.

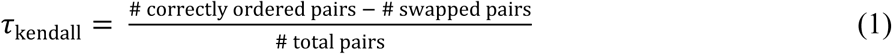

Various metrics summarize the accuracy with which the model fits the input data (Fig. 1B-E). The SVM^rank^ training metric shows varying agreement for all groups (*i.e.,* τ_kendall_ > 0) (Fig. 1B). For individual genes, activity values normalized and averaged across trials were not directly used for the training procedure (see Methods 4a); yet one would anticipate that scores for each gene should broadly correlate with the expression average. Indeed, the observed normalized activities positively correlate with the score (dubbed IMProve score for <u>I</u>ntegral <u>M</u>embrane <u>P</u>rotein expression improvement) output by the model (Fig. 1C, *ρ* > 0). Since SVM^rank^ transforms raw expression levels within each plate to ranks before training, there is no expectation or guarantee that magnitude differences in expression level manifest in magnitude differences in score. As a result, Spearman’s *ρ*, a rank correlation coefficient describing the agreement between two ranked quantities, is better suited for quantifying correlation over more common metrics like the R^2^ of a regression and Pearson’s *r*.

For a more quantitative approach to assessing the IMProve model’s success within the training data, we turn to the Receiver Operating Characteristic (ROC). ROC curves quantify the tradeoff between true positive and false positive predictions across the numerical scores output from a predictor. This is a more reliable assessment of prediction than simply calculating accuracy and precision from a single, arbitrary score threshold ^42^. The figure of merit that quantifies a ROC curve is the Area Under the Curve (AUC). Given that the AUC for a perfect predictor corresponds to 100% and that of a random predictor is 50% (Fig. 1D, grey dashed line), an AUC greater than 50% indicates predictive performance of the model (percentage signs hereafter omitted) (see Methods 5 and ^42^). Here, the ROC framework will be used to quantitatively assess the ability of our model to predict the outcomes within the various datasets.

The training datasets are quantitative measures of activity requiring that an activity threshold be chosen that defines positive or negative outcomes. For example, ROC curves using two distinct activity thresholds, at the 25^th^ or 75^th^ percentile of highest expression, are plotted with their calculated AUC values (Fig. 1D). While both show that the model has predictive capacity, a more useful visualization would consider all possible activity thresholds. For this, the AUC value for every activity threshold is plotted showing that the model has predictive power regardless of an arbitrarily chosen expression threshold (Fig. 1E). In total, the analysis demonstrates that the model can rank expression outcomes across all proteins in the training set. Interestingly, for PhoA-tagged proteins the model is progressively less successful with increasing activity. The implications for this are discussed later (see Fig. 2G below).

**Fig. 2.**
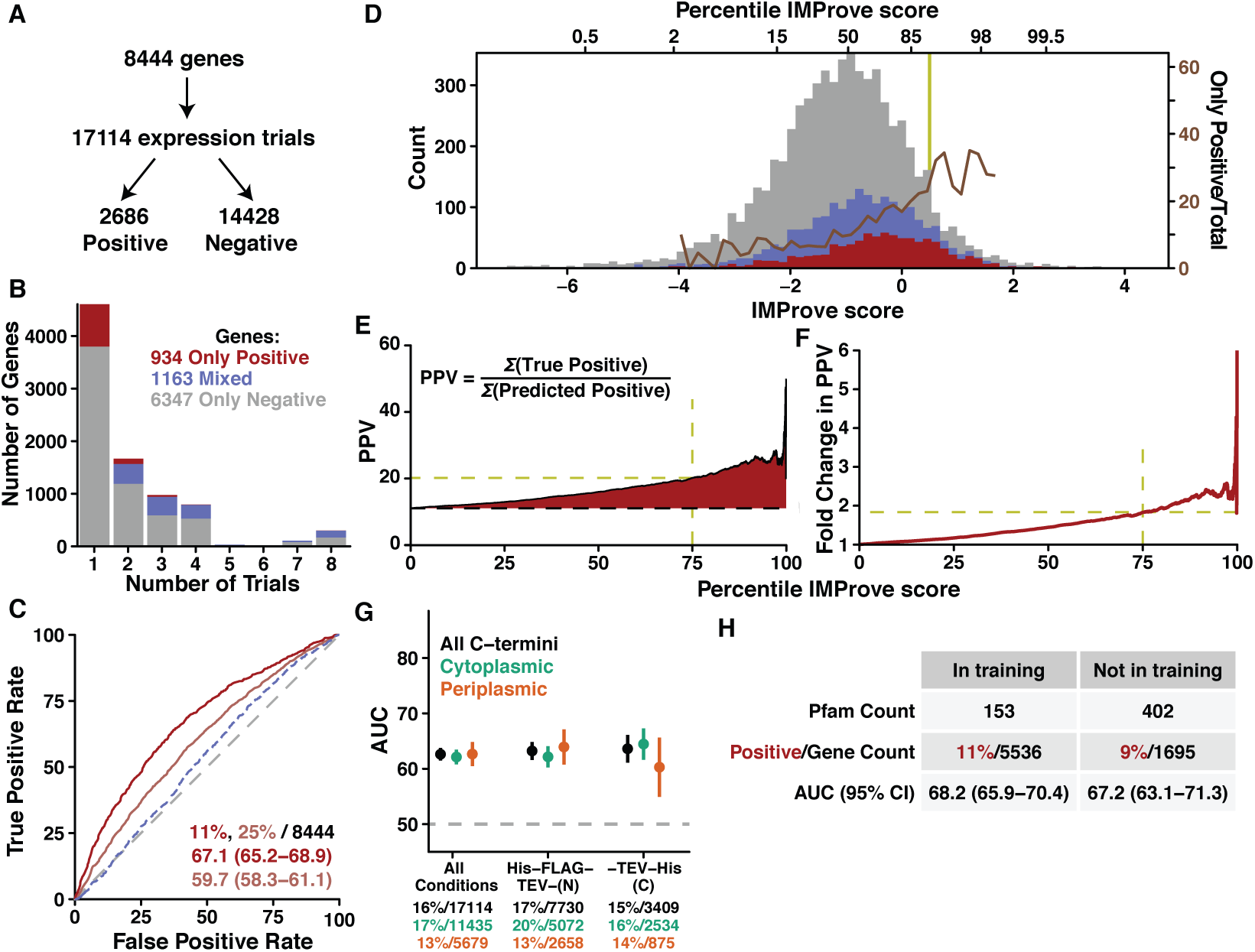
Success of the model against outcomes from NYCOMPS. **(A)** An overview of the NYCOMPS outcomes and **(B)** a histogram of the number of conditions tested per gene colored based on outcome. **(C)** Receiver Operating Characteristics for positive groupings given by Only Positive outcomes genes (red) and genes with at least one positive outcome (pink). The percent positive for each group (corresponding color), total counts (black), and Area Under the Curve (AUC) values with 95% Confidence Interval (CI) are shown. The ROC considering genes with Mixed outcomes only as positive is shown as a blue dashed line with an AUC of 53.5 (51.8-55.2). The grey dashed line shows the performance of a completely random predictor (AUC = 50). **(D)** Histograms of genes with Only Positive (red) and Only Negative outcomes (grey) across IMProve scores (binned as described in Methods 5). The percentage of Only Positive outcomes in each bin is overlaid as a brown line (right axis). **(E)** The Positive Predictive Value (PPV) plotted for each percentile IMProve score, *e.g.* 75 on the x-axis indicates the PPV for the top 25% of genes based on score for genes, where positive indicates genes with Only Positive outcomes. The dashed line shows the overall success rate of the NYCOMPS experimental outcomes (~11% Only Positive). **(F)** The fold change in the PPV as a function of IMProve score relative to the success rate of NYCOMPS. **(G)** The AUCs for outcomes across all trials and within the most-tested plasmids along with 95% CI. Performances are also split by predicted C-terminal localization ^39^. The numbers below indicate the total number of trials for each group and the percent within that group that were positive. **(H)** The NYCOMPS dataset split by the presence or absence of a Pfam family in the training set with AUCs calculated by considering Only Positive genes as positive outcomes.

### Demonstration of prediction against an independent large expression dataset

While the above analyses show that the model successfully fits the training data, we assess the broader applicability of the model outside the training set based on its success at predicting the outcomes of independent expression trials from distinct groups and across varying scales. The first test considers results from NYCOMPS, where 8444 IMP genes entered expression trials, in up to eight conditions, resulting in 17114 expression outcomes (Fig. 2A) ^2^. The majority of genes were attempted in only one condition (Fig. 2B), and, importantly, outcomes were non-quantitative (binary: expressed or not expressed) as indicated by the presence of a band by Coomassie staining of an SDS-PAGE gel after small-scale expression, solubilization, and nickel affinity purification ^3^. For this analysis, the experimental results are either summarized as outcomes per gene or broken down as raw outcomes across defined expression conditions. For outcomes per gene, we can consider various thresholds for considering a gene as positive based on NYCOMPS expression success (Fig. 2B). The most stringent threshold only regards a gene as positive if it has no negative outcomes (“Only Positive”, Fig. 2B, red). Since a well expressing gene would generally advance in the NYCOMPS pipeline without further small-scale expression trials, this positive group likely contains the best expressing proteins. A second category comprises genes with at least one positive and at least one negative trial (“Mixed”, Fig. 2B, blue). These genes likely include proteins that are more difficult to express.

ROCs assess predictive power across these groups (Fig. 2C). IMProve scores markedly distinguish genes in the most stringent positive group (Only Positive) from all other genes (AUC = 67.1) (Fig. 2C red). A permissive threshold considering genes as positive with at least one positive trial (Only Positive plus Mixed genes) shows moderate predictive power (Fig. 2C pink, AUC = 59.7). If instead the Mixed genes are considered alone (excluding the Only Positive), the model very weakly distinguishes the mixed group from Only Negative genes (Fig. 2C dashed blue, AUC = 53.5). This likely supports the notion that this pool largely consists of more difficult-to-express genes. For further analysis of NYCOMPS, we focus on the Only Positive pool as this likely represents the pool of best expressing proteins.

The predictive power of the IMProve model can be assessed by a variety of additional metrics. This can be qualitatively visualized as a histogram of the IMProve scores for genes separated by expression success (Only Positive, red; Mixed, blue; Only Negative, grey) (Fig. 2D). Visually, the distribution of the scores for the Only Positive group is shifted to a higher score relative to the Only Negative and Mixed groups. The dramatic increase in the percentage of Only Positive genes as a function of increasing IMProve score (overlaid as a brown line) further emphasizes this. A major aim of this work is to enrich the likelihood of choosing positively expressing proteins. The positive predictive value (PPV, true positives ÷ predicted positives) becomes a useful metric for positive enrichment as it conveys the degree of improved prediction over the experimental baseline of the dataset. The PPV of the model is plotted as a function of the percentile of the IMProve score for the Only Positive group (Fig. 2E). In the figure, the experimental baseline, all are predicted positive (PPV = 11.1%), is represented by a dashed line; therefore, a relative increase reflects the predictive power of the algorithm. For example, considering the top fourth of genes by IMProve score (75^th^ percentile, IMProve score = −0.2, PPV = 20%) shows that the algorithm increases the positive outcomes by 9% over baseline (1.82 fold change). Higher score cut-offs would have even higher increases in positive outcomes. For further illustration, we plot the fold-change in PPV across all thresholds (Fig. 2F).

We next confirm the ability of the IMProve model to predict within plasmids or sequence space distinct from those within the limited training set. For an overfit model, one might expect that only the subset of targets which most closely mirror the training data would show strong prediction. On the contrary, the model shows consistent performance throughout each of the eight distinct experimental conditions tested (Fig. 2G and Supplementary Table 2). One may also consider that the small size of the training set limited the number of protein folds sampled and, therefore, limited the number of folds that could be predicted by the model. To test this, we consider the performance of the model with regards to protein homology families, as defined by Pfam family classifications ^43^. The 8444 genes in the NYCOMPS dataset fall into 555 Pfam families (~15% not classified). To understand whether the IMProve score is biased towards families present in the training set, we separate genes in the NYCOMPS dataset into either within the 153 Pfam families found in the training set or outside this pool *(i.e.* not in the training set). Satisfyingly, there is no significant difference in AUC at 95% confidence between these groups (68.2 versus 67.2) (Fig. 2H). Combined, this highlights that the model is not sensitive to the experimental design of the training set and predicts broadly across different vector backbones and protein folds.

The ability to predict the experimental data from NYCOMPS allows returning to the question of alkaline phosphatase as a metric for expression. For the training set, proteins with C-termini in the periplasm show less consistent fitting by the model (Fig. 1, orange). To assess the generality of this result, the NYCOMPS outcomes are split into pools for either cytoplasmic or periplasmic C-terminal localization and AUCs are calculated for each. There are no significant differences in predictive capacity across all conditions (Fig. 2G, green vs. orange) irrespective of whether the tag is at the N- or C-terminus. This demonstrates that the IMProve model is applicable for all topologies.

At this point, it is useful to consider the potential improvement in the number of positive outcomes by using the IMProve model. NYCOMPS tested about a tenth of the 74 thousand possible IMPs from the 98 genomes of interest for expression ^2^. Had NYCOMPS tested the same number of genes from this pool, but selected to have an IMProve score greater than 0.5 (at the 91^st^ percentile (Fig. 2D, yellow line)), they would have increased their positive pool of 934 by an additional 1207 proteins. This represents a more than two-fold improvement in the return on investment and is a clear benchmark of success for the IMProve model.

### Further demonstration of prediction against small-scale independent datasets

The NYCOMPS example demonstrates the predictive power of the model across the broad range of sequence space encompassed by that dataset. Next, the performance of the model is tested against relevant subsets of sequence space *(e.g.* a family of proteins or the proteome from a single organism), which are reminiscent of laboratory-scale experiments that precede structural or biochemical analyses. While a number of datasets exist ^5,44–55^, we identified seven for which complete sequence information could be obtained to calculate all the necessary sequence features ^44–50^.

To understand the predictive performance within each of the small-scale datasets, we analyze the predictive performance of the model and highlight how the model could have been used to streamline those experiments. The clear predictive performance within the large-scale NYCOMPS dataset (Fig. 2) serves as a benchmark of expected performance at the scale of the experimental efforts for an individual lab (Fig. 3A). As targets within the various datasets were tested only one or a few times, experimental variability within each set could play a large-role on the outcomes noted. Therefore, we summarize positives within each dataset as those genes with the highest level of outcome as reported by the original authors as this outcome is likely most robust to such variability (*e.g*. seen via Coomassie Blue staining of an SDS-PAGE gel). To be complete, we have plotted and considered predictive performance across all possible outcomes as well (Fig. 3B-D, Supplementary Fig. 1).

**Fig. 3.**
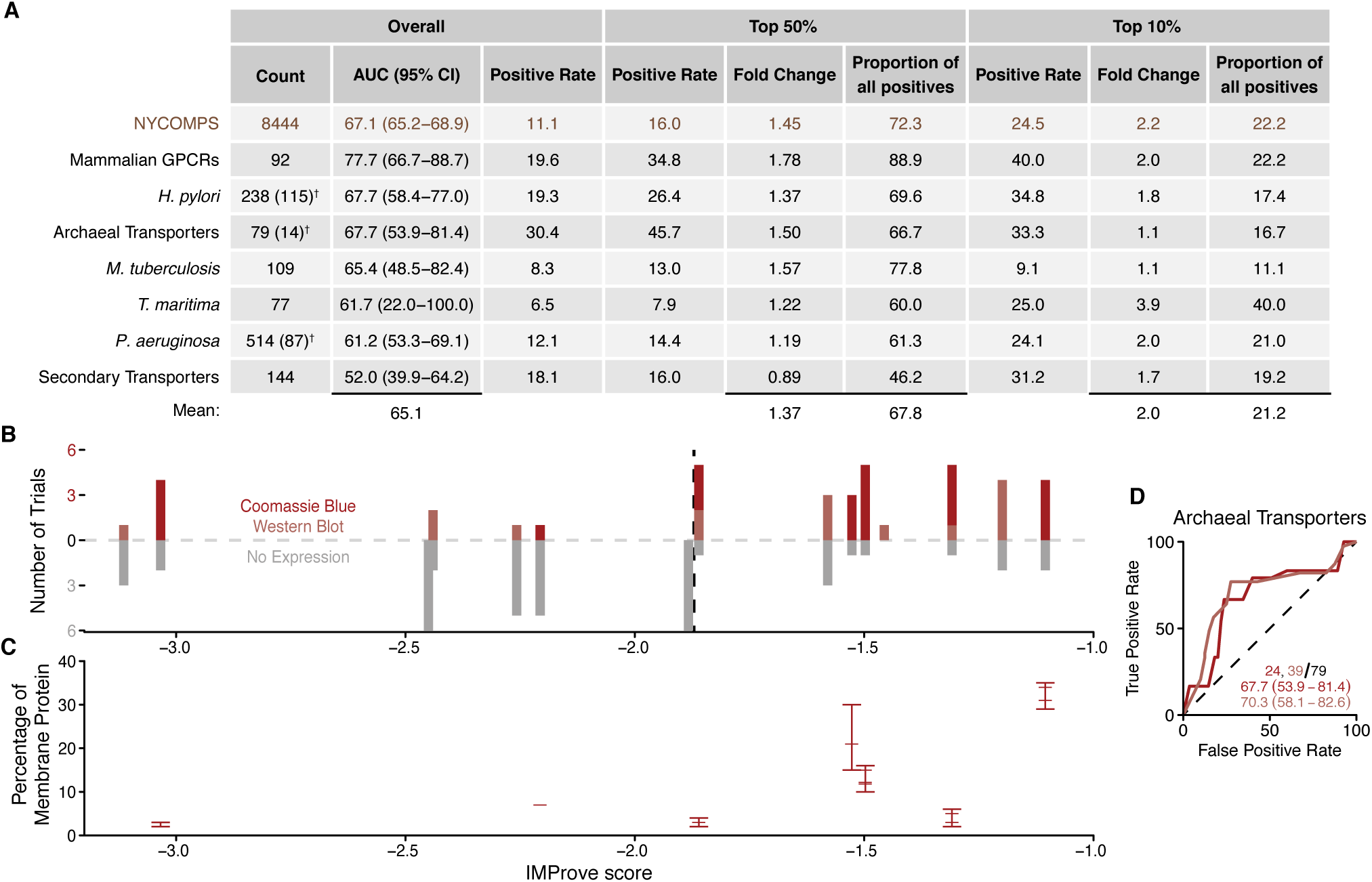
Success of the model against small scale outcomes. **(A)** Summary of the model’s performance against NYCOMPS and a variety of small scale expression experiments. Positive outcomes refer to those in the highest group as assigned by the authors of the respective studies. Where targets were tested in more than one condition (e.g. different plasmids or strains), the number of distinct proteins are indicated in parenthesis with a dagger. (**B**) The expression of archaeal transporters in up to 6 trials ^44^ Positive expression count is plotted above the dashed line and negative outcomes below the line. **(C)** Quantitative expression outcomes of those transporters as detected by Coomassie Blue. **(D)** Receiver Operating Characteristics (ROC) along with Areas Under the Curves (AUC) and 95% confidence interval as well as the total number of positives for the given threshold (red hues) along with the total outcomes (black) are presented. In each curve, increasing expression thresholds are displayed as deeper red.

The performance of the IMProve model for each of the small-scale datasets is consistent with that seen for the NYCOMPS dataset (Fig. 3A). This is most directly indicated by a mean AUC across all datasets of 65.6, highlighting the success across the varying scales. While the overall positive rate is different for each dataset, considering a cut-off in IMProve score, *e.g.* the top 50% or 10% of targets ranked by score, would have resulted in a greater percentage of positive outcomes. On average, ~70% of positives are captured within the top half of scores. Similarly, for the top 10% of scores, on average over 20% of the positives are captured. Simply put, for one tenth of the work one would capture a significant number of the positive outcomes within the pool of targets in each dataset.

For broader consideration, one can consider the fold change in positive rate by selecting targets informed by IMProve scores. Using the data available, only testing proteins within the top 10% of scores would result in an average fold change of 2.0 in the positive rate (*i.e*. twice as many positively expressed proteins). As positive rate is a bounded quantity (maximum is 100%), the possible fold change is bounded as well and becomes relative to the overall positive rate when considering various cut-offs (*e.g*. for *T. maritima* the maximum fold-change is 15.4 while for archaeal transporters it is 3.3). Taking the average maximum possible fold change (7.5), the IMProve model achieves nearly a third of the possible improvement in positive rate compared to a perfect predictor.

Since IMProve model was trained on quantitative expression outcomes, we also expect that it captures quantitative trends in expression, *i.e.* a higher score translates to greater amount of expressed protein. While the NYCOMPS results are consistent with this (Fig. 2b), of the various data sets, only the expression of archaeal transporters presents quantitative expression outcomes for consideration. For this dataset, 14 archaeal transporters were chosen based on their homology to human proteins ^44^ and tested for expression in *E. coli*; total protein was quantified in the membrane fraction by Coomassie Blue staining of an SDS-PAGE gel. Here, the majority of the expressing proteins fall into the higher half of the IMProve scores, 7 out of 9 of those with multiple positive outcomes (Fig. 3B). Strikingly, quantification of the Coomassie Blue staining highlights a clear correlation with the IMProve score where the higher expressing proteins have higher scores (Fig. 3C).

A final test considers the ability of the model to predict expression in hosts other than *E. coli*. The expression trials of 101 mammalian GPCRs in bacterial and eukaryotic systems ^47^ provides a data set for considering this question. For this experiment, trials in *E. coli* clearly follow the trend that IMProve can predict within this group of mammalian proteins (AUC = 77.7) (Fig. 3A & Supplementary Fig. 1A,B). However, the expression of the same set of proteins in *P. pastoris* fails to show any predictive performance (AUC = 54.8) (Supplementary Fig. 1A,B). This lack of predictive performance in *P. pastoris* suggests that the parameterization of the model, calibrated for *E. coli* expression, requires retraining to generate a different model that captures the distinct interplay of sequence parameters in other hosts.

### Biological importance of various sequence features

Considering the success of IMProve, one might anticipate that biological properties driving prediction may provide insight into IMP biogenesis and expression. Using a proof-of-concept linear model allowed for a straightforward and useful predictor. With a linear model, as employed here, extracting the importance of each feature is ordinarily straightforward; assuming features are distributed identically and independently (“i.i.d.”), the weight assigned to each feature should correspond to its relative importance. However, in our case, the input features do not satisfy these conditions and significant correlation exists between individual features (Supplementary Fig. 2). As a result, during the training procedure, unequal weight is placed across correlating features that represent the same underlying biological property, thereby, complicating the process of determining the biological underpinnings of the IMProve score. For example, the importance of transmembrane segment hydrophobicity for membrane partitioning is distributed between several features: among these the average ΔG_insertion_^38^ of TM segments has a positive weight whereas average hydrophobicity, a correlating feature, has a negative weight (Supplementary Table 1, Supplementary Fig. 2). As many features are correlated; conclusive information cannot be obtained simply using weights of individual features to interpret the relative importance of their underlying biological phenomena. We address this complication by coarsening our view of the features to two levels: First, we analyze features derived from protein versus those derived from nucleotide sequence, and then we look more closely at features groups after categorizing by biological phenomena.

The coarsest view of the features is a comparison of those derived from protein sequence versus those derived from nucleotide sequence. The summed weight for protein features is around zero, whereas for nucleotide features the summed weight is slightly positive suggesting that in comparison these features may be more important to the predictive performance of the model (Fig. 4A). Within the training set, protein features more completely explain the score both via correlation coefficients (Fig. 4B) as well as through ROC analysis (Fig. 4C). However, comparison of the predictive performance of the two subsets of weights shows that the nucleotide features alone can give similar performance to the full model for the NYCOMPS dataset (Fig. 4D). Within the small-scale datasets investigated, using only protein or nucleotide features shows no significant difference in predictive power at 95% confidence (Fig. 4E). In general, this suggests that neither protein nor nucleotide features are uniquely important for IMP expression. However, within the context of the trained model, nucleotide features are critical for predictive performance for a large and diverse dataset such as NYCOMPS. This finding corroborates growing literature that the nucleotide sequence holds significant determinants of biological processes full model for the NYCOMPS dataset (Fig. 4D). Within the small-scale datasets investigated, using only protein or nucleotide features shows no significant difference in predictive power at 95% confidence (Fig. 4E). In general, this suggests that neither protein nor nucleotide features are uniquely important for IMP expression. However, within the context of the trained model, nucleotide features are critical for predictive performance for a large and diverse dataset such as NYCOMPS. This finding corroborates growing literature that the nucleotide sequence holds significant determinants of biological processes ^40,26,56–58^

**Fig. 4.**
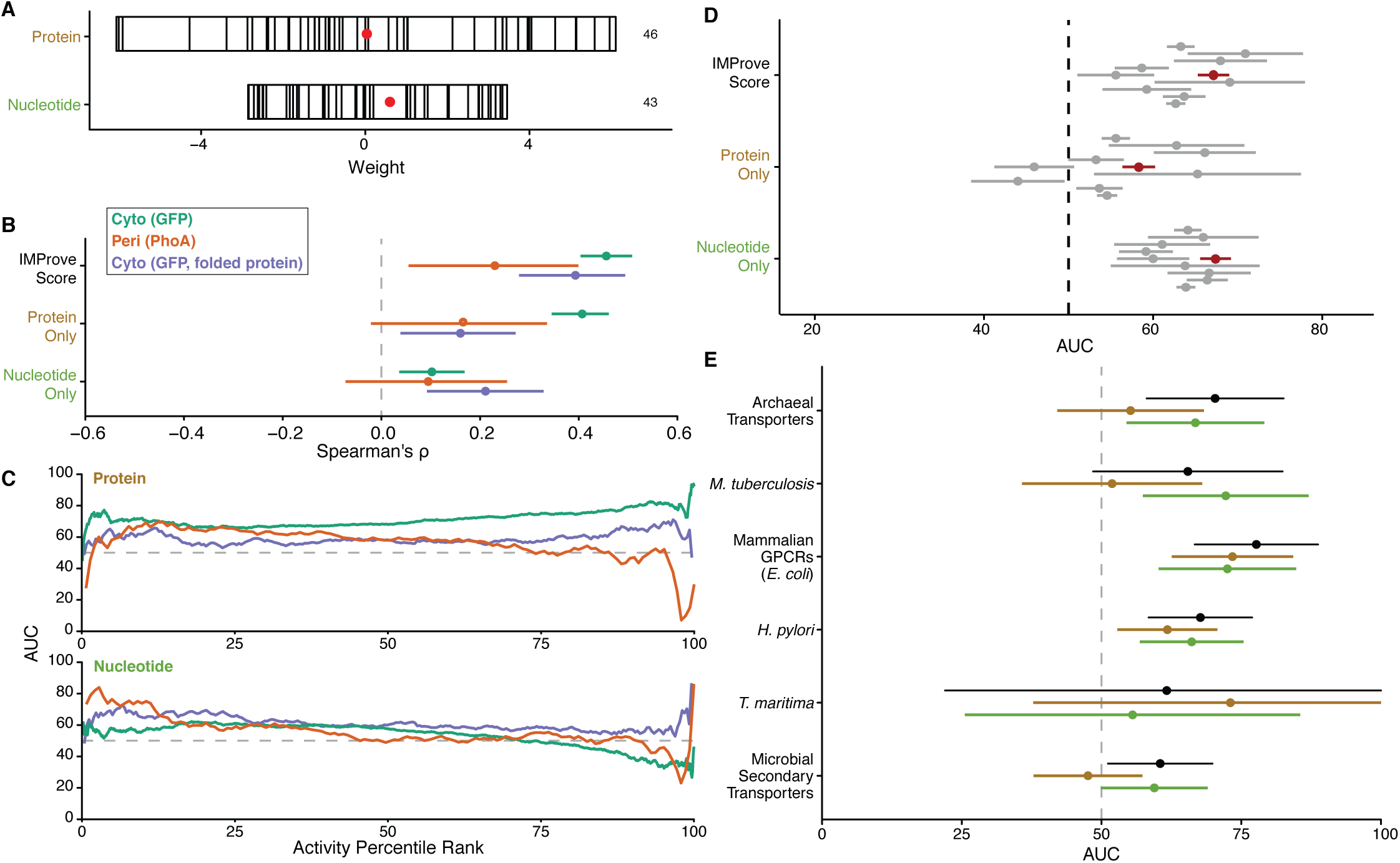
Feature contributions to the model. Classifying features by the type of sequence they are calculated from. **(B)** Considering the training set (as in Fig. 1), Spearman correlation coefficients with 95% confidence intervals using individual feature categories for each grouping of data within the training set of *E. coli* IMPs. Colors indicate the subset being assessed (green, whole cell GFP fluorescence; orange, alkaline phosphatase activity; purple, folded protein by in-gel fluorescence). **(C)** Protein/nucleotide feature dependence within the training set substantiated by the AUC of the ROC at every possible activity threshold for feature subsets independently (as in Fig. 1E). **(D)** The AUC and 95% confidence intervals using only protein or nucleotide features. **(E)** Protein/nucleotide feature dependence across small scale datasets shown as AUCs of the ROC along with 95% CI for the condition with the best overall predictive power (black).

We next collapse conceptually similar features into biological categories that allow us to infer the phenomena that drive prediction. Categories are chosen empirically *(e.g.* the hydrophobicity group incorporates sequence features such as average hydrophobicity, maximum hydrophobicity, ΔG_insertion_, *etc.*), which results in a reduction in overall correlation (Supplementary Fig. 3A). The full category list is provided in Supplementary Table 1. To visualize the importance of each category, the collapsed weights are summarized in Supplementary Fig. 3B, where each bar contains individual feature weights within a category. Features with a negative weight are stacked to the left of zero and those with a positive weight are stacked to the right. A red dot represents the sum of all weights, and the length of the bar gives the total absolute value of the combined weights within a category. Ranking the categories based on the sum of their weight suggests that some categories play a more prominent role than others. These include properties related to transmembrane segments (hydrophobicity and TM size/count), codon pair score, loop length, and overall length/pI.

To explore the role of each biological category in prediction, the performance of the model is assessed using only features within a given category. First, the strength of the correlation coefficients for given categories within the training set suggests the relative utility of each category for prediction. (Supplementary Fig. 3C, as in Fig. 4B). Examples of categories with high correlation coefficients are 5’ Codon Usage, Length/pI, Loop Length, and SD-like Sites. To verify their importance for prediction, we consider the AUC for prediction using each feature category for the NYCOMPS dataset (Supplementary Fig. 3D). In this analysis, only Length/pI shows some predictive power. Overall, the analysis of the training and large-scale testing dataset shows that no feature category independently drives the predictor. Excluding each individually does not significantly affect the overall predictive performance, except for Length/pI (data not shown). Sequence length composes the majority of the weight within this category and is one of the highest weighted features in the model (Supplementary Fig. 3A). This is consistent with the anecdotal observation that larger IMPs are typically harder to express. However, this parameter alone would not be useful for predicting within a smaller subset, like a single protein family, where there is little variance in length (*e.g*. Fig. 3,5). One might develop a predictor that was better for a given protein family under certain conditions with a subset of the entire features considered here; yet this would require *a priori* knowledge of the system, *i.e.* which sequence features were truly most important, and would preclude broad generalizability as shown for the IMProve model.

**Fig. 5.**
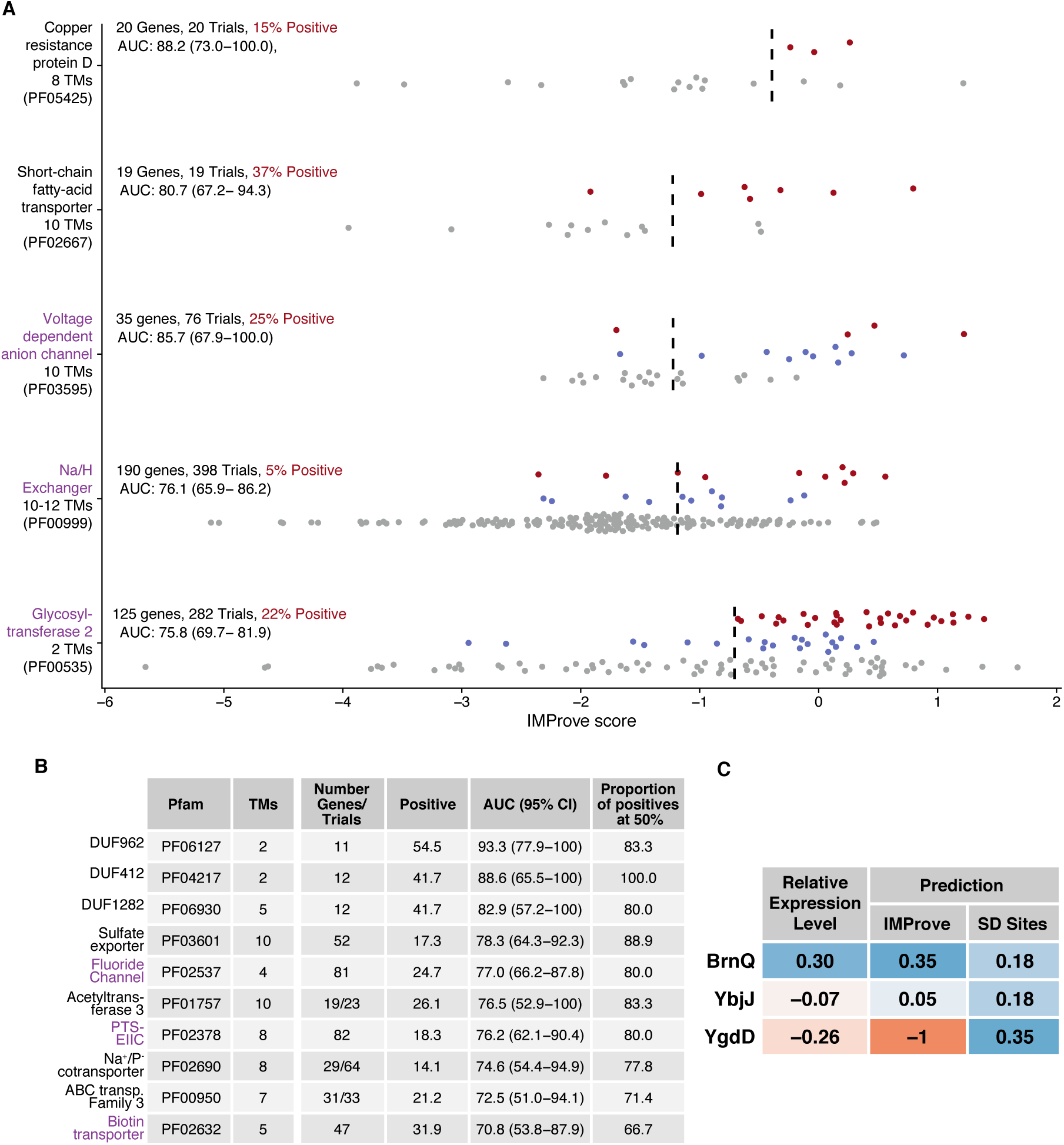
Usage of the model within IMP families and for optimization of expression. **(A)** Outcomes for specific protein families with an optimal IMProve score threshold indicated. Genes are shown in the chart as dots colored based on outcomes from trials: Only Positive (red), Only Negative (grey), and Mixed (blue). Overall statistics, as in Supplementary Table 3, are noted. Dashed lines represent the optimal threshold from the ROC curves. For the top two rows, each was only tested in a single condition (N: His-FLAG-TEV-gene). The bottom three rows are larger pools from NYCOMPS where there are multiple trials for many of the genes. **(B)** A table curated from Supplementary Table 3 where Pfams were selected based on specific criteria (minimum 10 trials, 4 positive and 4 negative outcomes) and ordered by AUC. Proteins, as in A, that have known crystal structures within the family are highlighted in purple. DUFs are domains of unknown function. For context, the following Pfam families correspond to TCDB classes: PF05425, <u>9.B.62</u>; PF02667, <u>2.A.73</u>; PF03595, <u>2.A.16</u>; PF00999, <u>2.A.36, 2.A.37</u>; PF00535, <u>9.B.32</u>; PF03601, <u>2.A.98</u>; PF02537, <u>1.A.43</u>; PF01757, <u>9.B.97</u>; PF02378, <u>4.A.1</u>, <u>4.A.2</u>, <u>4.A.3</u>; PF02690, <u>2.A.58</u>; PF02632, <u>2.A.88</u>^107^ **(C)** A comparison of the predictive capacity of IMProve compared to using silent mutations engineered to increase anti-SD sequence binding propensity ^26^. The table presents experimental relative expression level (mutant over wild-type sequence) versus predictions from relative changes in either IMProve score or SD-like sites. The cells are colored as a heat map from red (lower expression) to blue (higher expression).

### Usage of the IMProve model for IMP expression

We illustrate the IMProve model’s ability to identify promising homologs within a protein family by considering subsets of the broad range of targets tested by NYCOMPS. First, we consider two examples for protein families that do not have associated atomic resolution structures: copper resistance proteins (CopD, PF05425) and short-chain fatty-acid transporters (AtoE, PF02667). In the first two rows of Fig. 5A, genes from the two families are plotted by IMProve score and colored by experimental outcome. In both cases, as indicated by the AUCs of 88.2 and 80.7 (Fig. 5A), the model excels at predicting these families and provides a clear score cut-off to guide target selection for future expression experiments. For example, we expect that CopD homologs with IMProve scores above −1 will have a higher likelihood of expressing over other homologs.

We have calculated predictive performance for each Pfam found in the NYCOMPS data which allows us to provide considerations for future experiments (Supplementary Table 3). In particular, we highlight three families with many genes tested, multiple experimental trials and a spread of outcomes: voltage-dependent anion channels (PF03595), Na/H exchangers (PF00999), and glycosyltransferases (PF00535). For these, a very clear IMProve score cut-off emerges from the experimental outcomes (dashed line in Fig. 5A). Strikingly, for these families the IMProve model clearly ranks the targets with Only Positive outcomes (red) at higher scores, again suggesting a preference for the best expressing proteins (see Fig. 2 and 3). Similarly, many more families within NYCOMPS are predicted with high statistical confidence (Supplementary Table 3); we provide a subset as Fig. 5B. For these, if only genes in the top 50% of IMProve score were tested, 81% of the total positives would be captured. As noted before, this is a dramatic increase in efficiency. Excitingly, many of these families remain to be resolved structurally. Considering these results with the broader experimental data sets (Fig. 3), no matter the number of proteins one is willing to test, the IMProve model enables selecting targets with a high probability of expression success in *E. coli.*

### Sequence optimization for expression

The predictive performance of the model implies that the features defined here provide a coarse approximation of the fitness landscape for IMP expression. Attempting to optimize a single feature by modifying the sequence will likely affect the resulting score and expression due to changes in other features. Fluman, *et al.* provides an illustrative experiment ^26^. For that work, it was hypothesized that altering the number of Shine-Dalgarno (SD)-like sites in the coding sequence of a IMP would affect expression. To test this, silent mutations were engineered within the first 200 bases of three proteins (genes *ygdD, brnQ,* and *ybjJ* from *E. coli)* to increase the number of SD-like sites with the goal of improving expression. Expression trials demonstrated that only one of the proteins (BrnQ) had improved expression of folded protein. While the number of SD-like sites alone does not correlate, only 1 out of 3, the resulting changes in the IMProve score correlate with the changes in measured expression, 3 out of 3 (Fig. 5C). The IMProve model’s ability to capture the outcomes in this small test case illustrates the utility of integrating the contribution of the numerous parameters involved in IMP biogenesis.

## Discussion

Here, we have demonstrated a statistically driven predictor, IMProve, that decodes from sequence the sum of biological features that drive expression, a feat not previously possible ^10,17^. The current best practice for characterization of an IMP target begins with the identification and testing of multiple homologs or variants for expression. The predictive power of IMProve enables this by providing a low barrier-to-entry method to enrich more than two-fold the positive outcomes from such expression. IMProve allows for the prioritization of targets to test for expression making more optimal use of limited human and material resources. For groups with small scale projects such as individual labs, this means that for the same cost one would double the success rate. For large scale groups, such as companies or consortia, IMProve can reduce by half the cost required to obtain the same number of positive results. We provide the current predictor as a web service where scores can be calculated, and the method, associated data, and suggested analyses are publically available to catalyze progress across the community (clemonslab.caltech.edu).

Having shown that IMP expression can be predicted, the generalizability of the model is remarkable despite several known limitations. Using data from a single study for training precludes including certain variables that empirically influence expression such as the features corresponding to fusion tags and the context of the protein in an expression plasmid, *e.g.* the 5' untranslated region, for which there was no variation in the Daley, Rapp, *et al.* dataset. Moreover, using a simple proof-of-concept linear model allowed for a straightforward and robust predictor; however, intrinsically it cannot be directly related to the biological underpinnings. While we can extract some biological inference, a linear combination of sequence features does not explicitly reflect the reality of physical limits for host cells. To some extent, constraint information is likely encoded in the complex architecture of the underlying sequence space (e.g. through the genetic code, TM prediction, RNA secondary structure analyses). Future statistical models that improve on these limitations will likely hone predictive power and more intricately characterize the interplay of variables that underlie IMP expression in *E. coli* and other systems.

A perhaps surprising outcome of our results is the demonstration of the quantitatively important contribution of the nucleotide sequence as a component of the IMProve score. This echoes the growing literature that aspects of the nucleotide sequence are important determinants of protein biogenesis in general ^40,26,56–58^. While one expects that there may be different weights for various nucleotide derived features between soluble and IMPs, it is likely that these features are important for soluble proteins as well. An example of this is the importance of codon optimization for soluble protein expression, which has failed to show any general benefit for IMPs ^10^. Current expression predictors that have predictive power for soluble proteins have only used protein sequence for deriving the underlying feature set ^59,60^. Future prediction methods will likely benefit from including nucleotide sequence features as done here.

The ability to predict phenotypic results using sequence based statistical models opens a variety of opportunities. As done here, this requires a careful understanding of the system and its underlying biological processes enumerated in a multitude of individual variables that impact the stated goal of the predictor, in this case enriching protein expression. As new features related to expression are discovered, future work will incorporate these leading to improved models. This can include features derived from other approaches such as the integration efficiency derived from coarse-grained molecular dynamics ^12,15^. Based on these results, expanding to new expression hosts such as eukaryotes seems entirely feasible, although a number of new features may need to be considered, *e.g.* glycosylation sites and trafficking signals. Moreover, the ability to score proteins for expressibility creates new avenues to computationally engineer IMPs for expression. The proof-of-concept described here required significant work to compile data from genomics consortia and the literature in a readily useable form. As data becomes more easily accessible, broadly leveraging diverse experimental outcomes to decode sequence-level information, an extension of this work, is anticipated.

## Author Contributions

S.M.S., A.M., and W.M.C. conceived the project. S.M.S. developed the approach. S.M.S., A.M., and N.J. compiled sequence and experimental data. N.J. created code to demonstrate feasibility. S.M.S. performed all published calculations. S.M.S. and W.M.C. wrote the manuscript.

## Acknowledgements

We thank Daniel Daley and Thomas Miller’s group for discussion, Yaser Abu-Mostafa and Yisong Yue for guidance regarding machine learning, Niles Pierce for providing NUPACK source code ^30^, Welison Floriano and Naveed Near-Ansari for maintaining local computing resources, and Samuel Schulte for suggesting the model’s name. We thank Michiel Niesen, Stephen Marshall, Thomas Miller, Reid van Lehn, James Bowie, and Tom Rapoport for comments on the manuscript. Models and analyses are possible thanks to raw experimental data provided by Daniel Daley and Mikaela Rapp ^16^; Nir Fluman ^26^; Edda Kloppmann, Brian Kloss, and Marco Punta from NYCOMPS ^2,3^. Pikyee Ma ^44^; Renaud Wagner ^47^; Florent Bernaudat ^51^, and Constance Jeffrey ^45^.

We acknowledge funding from an NIH Pioneer Award to WMC (5DP1GM105385); a Benjamin M. Rosen graduate fellowship, a NIH/NRSA training grant (5T32GM07616), and a NSF Graduate Research fellowship to SMS; and an Arthur A. Noyes Summer Undergraduate Research Fellowship to NJ. Computational time was provided by Stephen Mayo and Douglas Rees. This material is based upon work supported by the National Science Foundation Graduate Research Fellowship Program under Grant No. 1144469. Any opinions, findings, and conclusions or recommendations expressed in this material are those of the authors and do not necessarily reflect the views of the National Science Foundation. This work used the Extreme Science and Engineering Discovery Environment (XSEDE), which is supported by National Science Foundation grant number ACI-1053575 ^61^.

## Online Methods

Sequence mapping & retrieval and feature calculation was performed in Python 2.7 ^62^ using BioPython ^63^ and NumPy ^64^; executed and consolidated using Bash (shell) scripts; and parallelized where possible using GNU Parallel ^65^. Data analysis and presentation was done in R ^66^ within RStudio ^67^ using magrittr ^68^, plyr ^69^, dplyr ^70^, asbio ^71^, and datamart ^72^ for data handling; ggplot2 ^73^, ggbeeswarm ^74^, GGally ^75^, gridExtra ^76^, cowplot ^77^, scales ^78^, viridis ^79^, and RColorBrewer ^80,81^ for plotting; multidplyr ^82^ with parallel ^66^ and foreach ^83^ with iterators ^84^ and doMC ^85^/doParallel ^86^ for parallel processing; and roxygen2 ^87^ for code organization and documentation as well as other packages as referenced.

### 1. Collection of data necessary for learning and evaluation

#### *E. coli* Sequence Data

The nucleotide sequences from ^16^ were deduced by reconstructing forward and reverse primers *(i.e.* ~20 nucleotide stretches) from each gene in Colibri (based on EcoGene 11), the original source cited and later verified these primers against an archival spreadsheet provided directly by Daniel Daley (personal communication). To account for sequence and annotation corrections made to the genome after Daley, Rapp, *et al.’s* work, these primers were directly used to reconstruct the amplified product from the most recent release of the *E. coli* K-12 substr. MG1655 genome ^35^ (EcoGene 3.0; U00096.3). Although Daniel Daley mentioned that raw reads from the Sanger sequencing runs may be available within his own archives, it was decided that the additional labor to retrieve this data and parse these reads would not significantly impact the model. The deduced nucleotide sequences were verified against the protein lengths given in Supplementary Table 1 from ^16^. The plasmid library tested in ^26^ was provided by Daniel Daley, and those sequences are taken to be the same.

#### *E. coli* Training Data

The preliminary results using the mean-normalized activities echoed the findings of ^16^ that these do not correlate with sequence features either in the univariate sense (many simple linear regressions, Supplementary Table 1 ^16^) or a multivariate sense (multiple linear regression, data not shown). This is presumably due to the loss of information regarding variability in expression level for given genes or due to the increase in variance of the normalized quantity (See Methods 4a) due to the normalization and averaging procedure. Daniel Daley and Mikaela Rapp provided spreadsheets of the outcomes from the 96-well plates used for their expression trials and sent scanned copies of the readouts from archival laboratory notebooks where the digital data was no longer accessible (personal communication). Those proteins without a reliable C-terminal localization (as given in the original work) or without raw expression outcomes were not included in further analyses.

Similarly, Nir Fluman also provided spreadsheets of the raw data from the set of three expression trials performed in ^26^.

#### New York Consortium on Membrane Protein Structure (NYCOMPS) Data

Brian Kloss, Marco Punta, and Edda Kloppman provided a dataset of actions performed by the NYCOMPS center including expression outcomes in various conditions ^2,3^. The protein sequences were mapped to NCBI GenInfo Identifier (GI) numbers either via the Entrez system ^88^ or the Uniprot mapping service^89^. Each GI number was mapped to its nucleotide sequence via a combination of the NCBI Elink mapping service and the “coded_by” or “locus” tags of Coding Sequence (CDS) features within GenBank entries. Though a custom script was created, a script from Peter Cock on the BioPython listserv to do the same task via a similar mapping mechanism was found ^90^. To confirm all the sequences, the TargetTrack ^23^ XML file was parsed for the internal NYCOMPS identifiers and compared for sequence identity to those that had been mapped using the custom script; 20 (less than 1%) of the sequences had minor inconsistencies and were manually replaced.

#### Archaeal transporters Data

The locus tags (“Gene Name” in Table 1) were mapped directly to the sequences and retrieved from NCBI ^44^. Pikyee Ma and Margarida Archer clarified questions regarding their work to inform the analysis.

#### GPCR Expression Data

Nucleotide sequences were collected by mapping the protein identifiers given in Table 1 from ^47^ to protein GIs via the Uniprot mapping service ^89^ and subsequently to their nucleotide sequences via the custom mapping script described above (see NYCOMPS). The sequence length and pI were validated against those provided. Renaud Wagner assisted in providing the nucleotide sequences for genes whose listed identifiers were unable to be mapped and/or did not pass the validation criteria as the MeProtDB (the sponsor of the GPCR project) does not provide a public archive.

#### *Helicobacter pylori* Data

Nucleotide sequences were retrieved by mapping the locus tags given in Supplemental Table 1 from ^48^ to locus tags in the Jan 31, 2014 release of the *H. pylori* 26695 genome (AE000511.1). To verify sequence accuracy, sequences whose molecular weight matched that given by the authors were accepted. Those that did not match, in addition to the one locus tag that could not be mapped to the Jan 31, 2014 genome version, were retrieved from the Apr 9, 2015 release of the genome (NC_000915.1). Both releases are derived from the original sequencing project ^91^. After this curation, all mapped sequences matched the reported molecular weight.

In this data set, expression tests were performed in three expression vectors and scored as 1, 2, or 3. Two vectors were scored via two methods. For these two vectors, the two scores were averaged to give a single number for the condition making them comparable to the third vector while yielding 2 additional thresholds (1.5 and 2.5) result in the 5 total curves shown (Supplementary Fig. 2B).

#### *Mycobacterium tuberculosis* Data

The authors note using TubercuList through GenoList ^92^, therefore, nucleotide sequences were retrieved from the archival website based on the original sequencing project ^93^. The sequences corresponding to the identifiers and outcomes in Table 1 from ^46^ were validated against the provided molecular weight.

#### *Secondary Transporter* Data

GI Numbers given in Table 1 from ^50^ were matched to their CDS entries using the custom mapping script described above (see NYCOMPS). Only expression in *E. coli* with IPTG-inducible vectors was considered.

#### *Thermotoga maratima* Data

Gene names given in Table 1 ^94^ were matched to CDS entries in the Jan 31, 2014 release of the *Thermotoga maritima* MSB8 genome (AE000512.1), a revised annotation of the original release ^95^. The sequence length and molecular weight were validated against those provided.

#### *Pseudomonas aeruginosa* Data

Outcomes in Additional file 1 ^45^ were matched to coding sequences provided by Constance Jeffrey.

#### Shine-Dalgarno-like mutagenesis Data

Folded protein is quantified by densitometry measurement ^96,97^ of the relevant band in Figure 6 of ^26^. Relative difference is calculated as is standard:

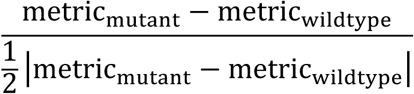

### 2. Details related to the calculation of sequence features

Transmembrane segment topology was predicted using Phobius Constrained for the training data and Phobius for all other datasets ^39^. We were able to obtain the Phobius code and integrate it directly into our feature calculation pipeline resulting in significantly faster speeds than any other option. Several features were obtained by averaging per-site metrics (e.g. per-residue RONN3.2 disorder predictions) in windows of a specified length. Windowed tAI metrics are calculated over *all* 30 base windows (not solely over 10 codon windows). Supplementary Table 1 includes an in-depth description of each feature. Future work will explore contributions of elements outside the gene of interest, *e.g.* ribosomal binding site, linkers, tags.

### 3. Preparation for model learning

Calculated sequence features for the IMPs in the *E. coli* dataset as well as raw activity measurements, *i.e.* each 96-well plate, were loaded into R. As is best practice in using Support Vector Machines, each feature was “centered” and “scaled” where the mean value of a given feature was subtracted from each data point and then divided by the standard deviation of that feature using `preprocess` ^98^. As is standard practice, the resulting set was then culled for those features of near zero-variance, over 95% correlation (Pearson’s *r*), and linear dependence (`nearZeroVar`, `findCorrelation`, `findLinearCombos`)^98^. In particular this procedure removed extraneous degrees of freedom during the training process which carry little to no additional information with respect to the feature space and which may over represent certain redundant features. Features and outcomes for each list (“query”) were written into the SVM^light^ format using a modified `svmlight.write` ^99^.

The final features were calculated for each sequence in the test datasets, prepared for scoring by “centering” and “scaling” by the training set parameters via `preprocess` ^98^, and then written into SVM^light^ format again using a modified `svmlight.write`.

### 4. Model selection, training, and evaluation using SVM^rank^

**a.** At the most basic level, our predictive model is a learned function that maps the parameter space (consisting of nucleotide and protein sequence features) to a response variable (expression level) through a set of governing weights (*w*_1_, *w*_2_,…, *w*_*N*_). Depending on how the response variable is defined, these weights can be approximated using several different methods. As such, defining a response variable that is reflective of the available training data is key to selecting an appropriate learning algorithm.

The quantitative 96-well plate results ^16^ that comprise our training data do not offer an absolute expression metric valid over all plates—the top expressing proteins in one plate would not necessarily be the best expressing within another. As such, this problem is suited for preference-ranking methods. As a ranking problem, the response variable is the ordinal rank for each protein derived from its overexpression relative to the other members of the same plate of expression trials. In other words, the aim is to rank highly expressed proteins (based on numerous trials) at higher scores than lower expressed proteins by fitting against the <u>order</u> of expression outcomes from each constituent 96-well plate.

**b.** As the first work of this kind, the aim was to employ the simplest framework necessary taking in account the considerations above. The method chosen computes all valid pairwise classifications (*i.e*. within a single plate) transforming the original ranking problem into a binary classification problem. The algorithm outputs a score for each input by minimizing the number of swapped pairs thereby maximizing Kendall’s τ ^100^. For example, consider the following data generated via context A (*X*_A,1_,*Y*_A,1_), (*X*_A,2_, *Y*_A,2_) and B (*X*_B,1_,*Y*_B,1_),(*X*_B,2_,*Y*_B,2_) where observed response follows as index *i, i.e. Y*_n_< *Y*_n+1_. Binary classifier *f* (*X*_*i*_,*X*_*j*_) gives a score of 1 if an input pair matches its ordering criteria and − 1 if not, *i.e. Y*_*i*_ < *Y*_*j*_:

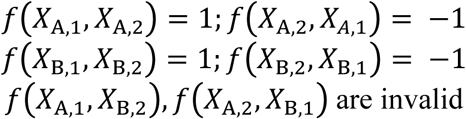

Free parameters describing *f* are calculated such that those calculated orderings *f*(*X*_A,1_), *f* (*X*_A,2_)…; *f*(*X*_B,1_), *f*(*X*_B,2_)…most closely agree (overall Kendall’s *τ*) with the observed ordering *Y*_*n*_, *Y*_n+1_,…. In this sense, *f* is a pairwise Learning to Rank method.

Within this class of models, a linear preference-ranking Support Vector Machine was employed ^101^. To be clear, as an algorithm a preference-ranking SVM operates similarly to the canonical SVM binary classifier. In the traditional binary classification problem, a linear SVM seeks the maximally separating hyper-plane in the feature space between two classes, where class membership is determined by which side of the hyper-plane points reside. For some *n* linear separable training examples *D* = {(*x*_*i*_)|*x*_*i*_ ϵ ℝ^*d*^}^*n*^ and two classes *y*_*i*_ ϵ {−1,1}, a linear SVM seeks a mapping from the *d*-dimensional feature space ℝ^*d*^ → {-1,1} by finding two maximally separated hyperplanes *w* ⋅ *x* − *b* = 1 and *w* ⋅ *x* − *b* = − 1 with constraints that *w* ⋅ *x*_*i*_ − *b* ≥ 1 for all *x*_*i*_ with *y*_*i*_ *ϵ*{1} and *w* ⋅ *x*_*i*_ − *b* ≤ − 1 for all *x*_*i*_ with *y*_*i*_ *ϵ* {−1}. The feature weights correspond to the vector *w*, which is the vector perpendicular to the separating hyperplanes, and are computable in O(*n* log *n*) implemented as part of the SVM^rank^ software package, though in O(*n*^2^) ^41^. See ^101^ for an in-depth, technical discussion.

**c.** In a soft-margin SVM where training data is not linearly separable, a tradeoff between misclassified inputs and separation from the hyperplane must be specified. This parameter *C* was found by training models against raw data from Daley, Rapp, *et al.* with a grid of candidate *C* values (2^*n*^∀ *n ϵ* [−5, 5]) and then evaluated against the raw “folded protein” measurements from Fluman, *et al.* The final model was chosen by selecting that with the lowest error from the process above (*C* = 2^5^). To be clear, the final model is composed solely of a single weight for each feature; the tradeoff parameter *C* is only part of the training process.

Qualitatively, such a preference-ranking method constructs a model that ranks groups of proteins with higher expression level higher than other groups with lower expression value. In comparison to methods such as linear regression and binary classification, this approach is more robust and less affected by the inherent stochasticity of the training data.

### 5. Quantitative Assessment of Predictive Performance

In generating a predictive model, one aims to enrich for positive outcomes while ensuring they do not come at the cost of increased false positive diagnoses. This is formalized in Receiver Operating Characteristic (ROC) theory (for a primer see ^42^), where the true positive rate is plotted against the false positive rate for all classification thresholds (score cutoffs in the ranked list). In this framework, the overall ability of the model to resolve positive from negative outcomes is evaluated by analyzing the Area Under a ROC curve (AUC) where AUC_perfect_=100% and AUC_random_=50% (percentage signs are omitted throughout the text and figures). All ROCs are calculated through pROC ^102^ using the analytic Delong method for AUC confidence intervals ^103^. Bootstrapped AUC CIs (N = 10^6^) were precise to 4 decimal places suggesting that analytic CIs are valid for the NYCOMPS dataset.

With several of our datasets, no definitive standard or clear-cut classification for positive expression exists. However, the aim is to show and test all reasonable classification thresholds of positive expression for each dataset in order to evaluate predictive performance as follows:

#### Training data

The outcomes are quantitative (activity level), so each ROC is calculated by normalizing within each dataset to the standard well subject to the discussion in 4a above (LepB for PhoA, and InvLepB for GFP) (examples in Fig. 1D) for each possible threshold, *i.e.* each normalized expression value with each AUC plotted in Fig. 1E. 95% confidence intervals of Spearman’s ρ are given by 10^6^ iterations of a bias-corrected and accelerated (BCa) bootstrap of the data (Fig. 1A,C) ^104^ Large-scale - ROCs were calculated for each of the expression classes (Fig. 2E). Regardless of the split, predictive performance is noted. The binwidth for the histogram was determined using the Freedman-Diaconis rule^105^, and scores outside the plotted range comprising <0.6% of the density were implicitly hidden.

#### Large-scale

ROCs were calculated for each of the expression classes (Fig. 2E). Regardless of the split, predictive performance is noted. The binwidth for the histogram was determined using the Freedman-Diaconis rule^105^, and scores outside the plotted range comprising <0.6% of the density were implicitly hidden.

#### Small-scale

Classes can be defined in many different ways. To be principled about the matter, ROCs for each possible cutoff are presented based on definitions from each publication (Fig. 3C,E,G, Supplementary Fig. 2B,D,F). See Methods 1 for any necessary details about outcome classifications for each dataset.

### 6. Feature Weights

Weights for the learned SVM are pulled directly from the model file produced by SVM^light^ and are given in Supplementary Table 1.

### 8. Availability

All analysis is documented in a series of R notebooks ^106^ available openly at github.com/clemlab/IMProve. These notebooks provide fully executable instructions for the reproduction of the analyses and the generation of figures and statistics in this study. The IMProve model is available as a web service at clemonslab.caltech.edu. Additional code is available upon request.

## Supplementary Material

**Supplementary Fig. 1.**
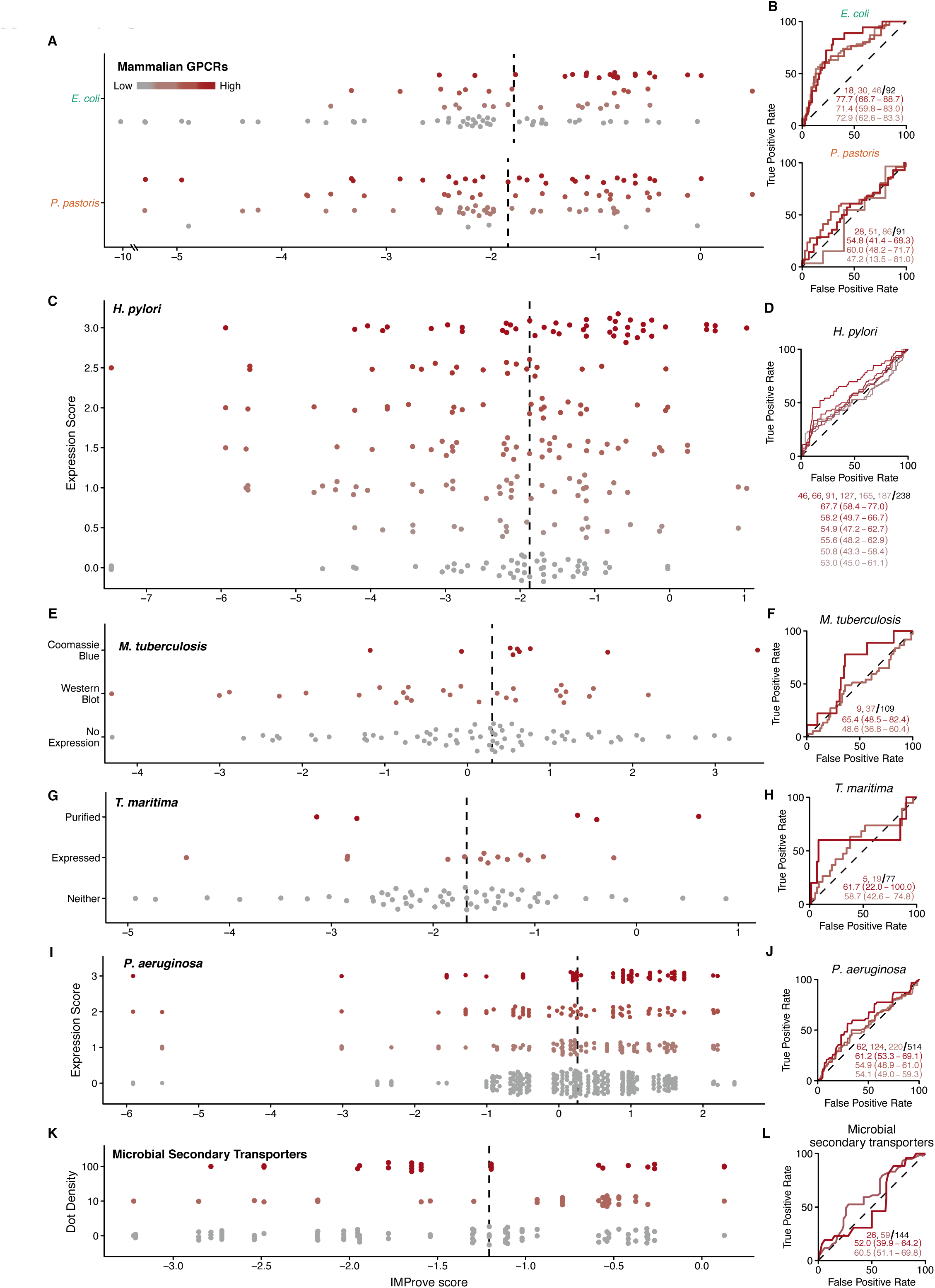
Success of the model against a variety of small scale outcomes. For each set, vertical lines indicate the median IMProve score. Receiver Operating Characteristics (ROC) along with Areas Under the Curves (AUC) and 95% confidence interval as well as the total number of positives for the given threshold (red hues) along with the total outcomes (black) are presented. In each curve, increasing expression thresholds as defined by the original publication are displayed as deeper red. The Reciever Operating Characteristic (ROC) with each cutoff is plotted, where a higher cutoff is represented by a deeper red, followed by the Area Under the Curves (directly below) in colors that correspond to the respective curve. **(A,B)** Mammalian GPCR expression in either *E. coli* (top) or *P. pastoris* (bottom). **(C,D)** Experimental expression of 116 *H. pylori* membrane proteins in *E. coli* in at most 3 vectors (238 trials) scored as either a 1, 2, or 3 from the outcome of a dot blot as well as Coomassie Staining of an SDS-PAGE gel for two of the vectors. To compare the three vectors with a single set of scores, the two scores were averaged to give a single number for a condition making them comparable to the third vector while yielding 2 additional thresholds (1.5 and 2.5) and the 6 total levels shown. **(E,F)** Experimental expression of *M. tuberculosis* membrane proteins plotted based on outcomes. **(G,H)** Pooled outcomes from the expression of 87 *P. aeruginosa* membrane proteins in *E. coli* across 3 plasmids and 2 strains scored on a relative scale. **(I,J)** Expression of 77 *T. maritima* membrane proteins in *E. coli* noted as purified (5), not purified but expressed (14), or neither. **(K,L)** Expression of 37 microbial secondary transporters in 4 IPTG-inducible vectors (144 trials) in*E. coli* quantified as 10 ng/mL (pink) or 100 ng/mL (red) via dot blot.

**Supplementary Fig. 2.**
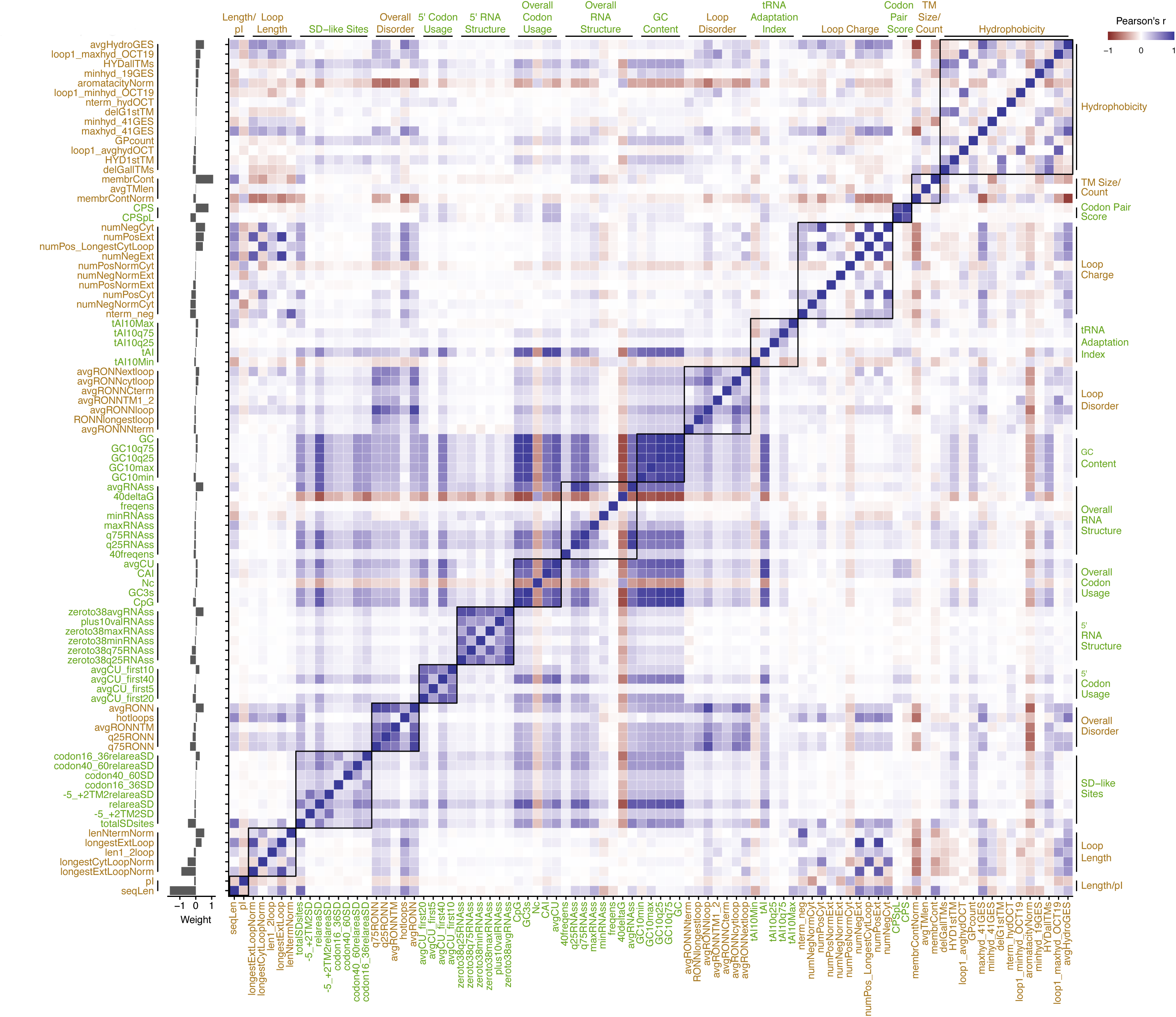
Complete set of feature correlations and their individual contributions to the model. Features are ordered first by category and then by weight (grey bars). Labels are green for protein-sequence derived and brown for nucleotide-sequence derived features. Pearson correlation coefficient between each pair of features across the NYCOMPS dataset is plotted (right). See S1 Table for a detailed description of each feature. Feature categories are overlaid as square boxes and indicated by black bars on the top, left, and right of the correlation matrix.

**Supplementary Fig. 3.**
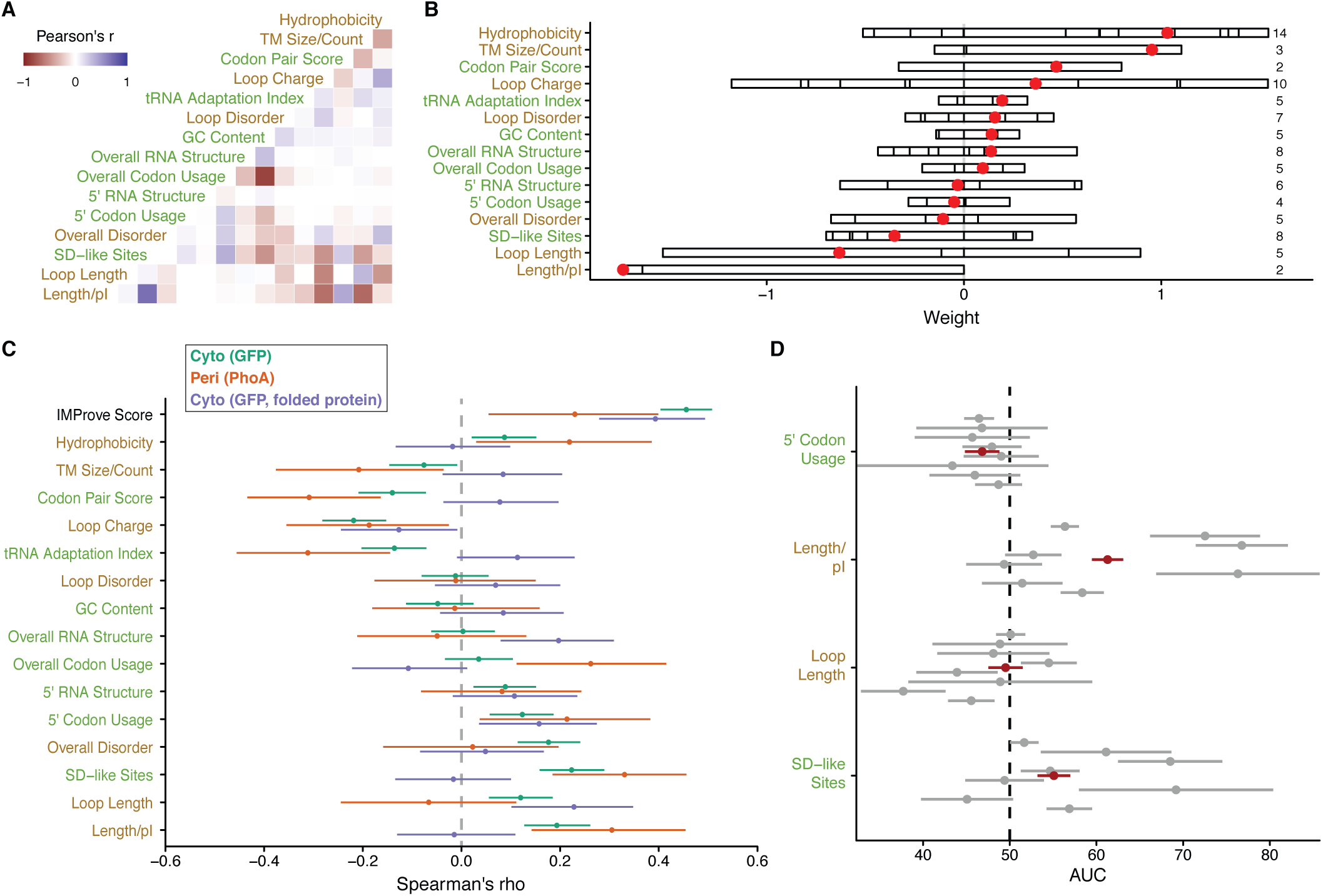
Feature contributions to the model across datasets used for training and validation. **(A)** Pearson correlation coefficients between feature categories are shown. Feature labels are green for protein-sequence derived and brown for nucleotide-sequence derived. **(B)** Total weight for each category is represented as a bar. The contribution of each feature to the category is shown by partitioning the bar. The red dot indicates the total sum of weights within the category. **(C)** Feature category dependence within the training set is shown by Spearman’s ρ and 95% CI between the normalized outcomes versus the feature subset. **(D)** Considering the NYCOMPS data set (as in Fig 2), the Area Under the Curve (AUC) of a Receiver Operating Characteristic and 95% confidence interval when predicting solely by features from the specified category against the NYCOMPS dataset. Red, using positive only as the cut-off for individual genes (Fig 2C); grey, using positive outcomes within each plasmid and solubilization condition (as in Fig 2E).

**Supplementary Table 1.**
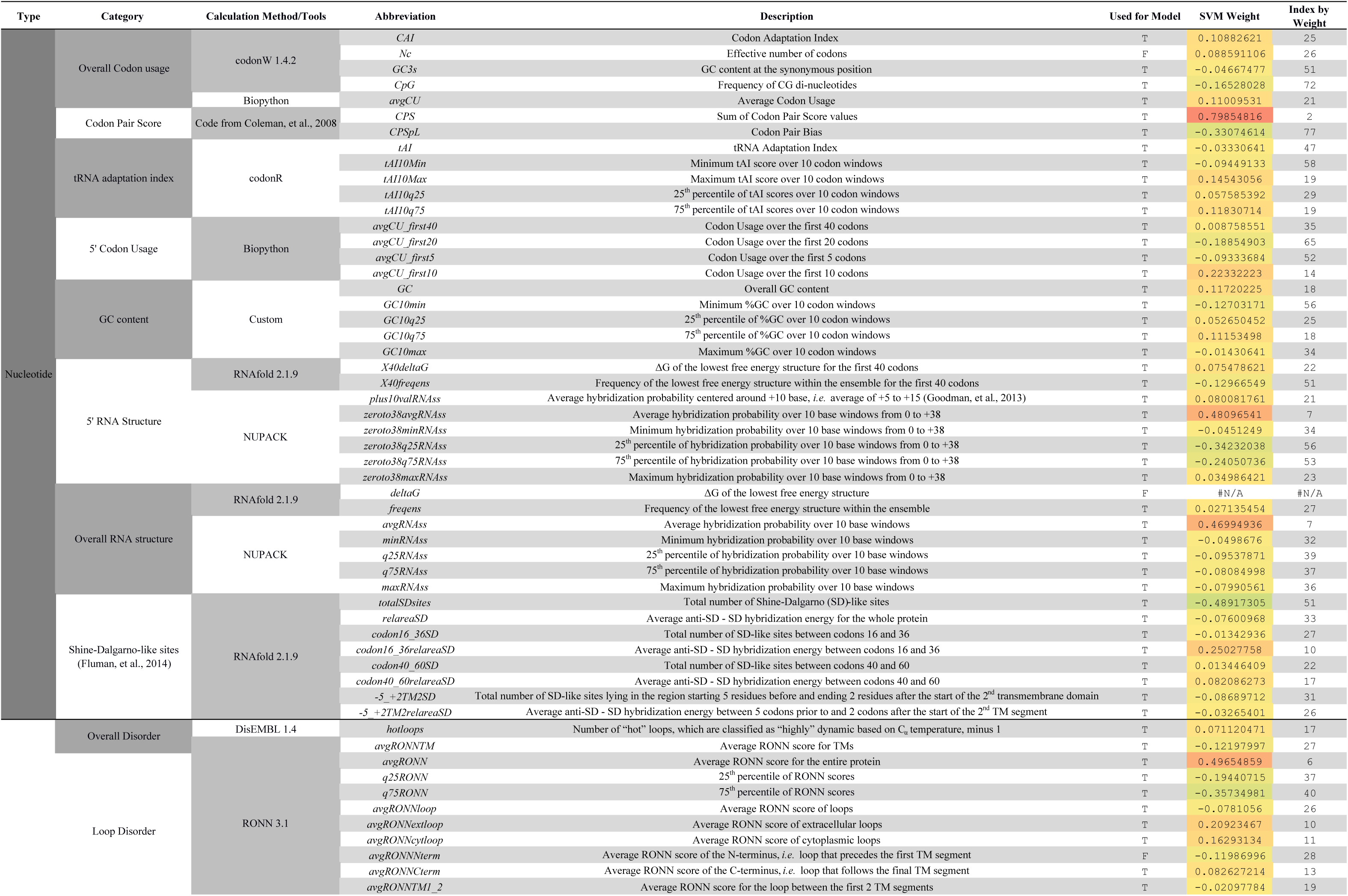
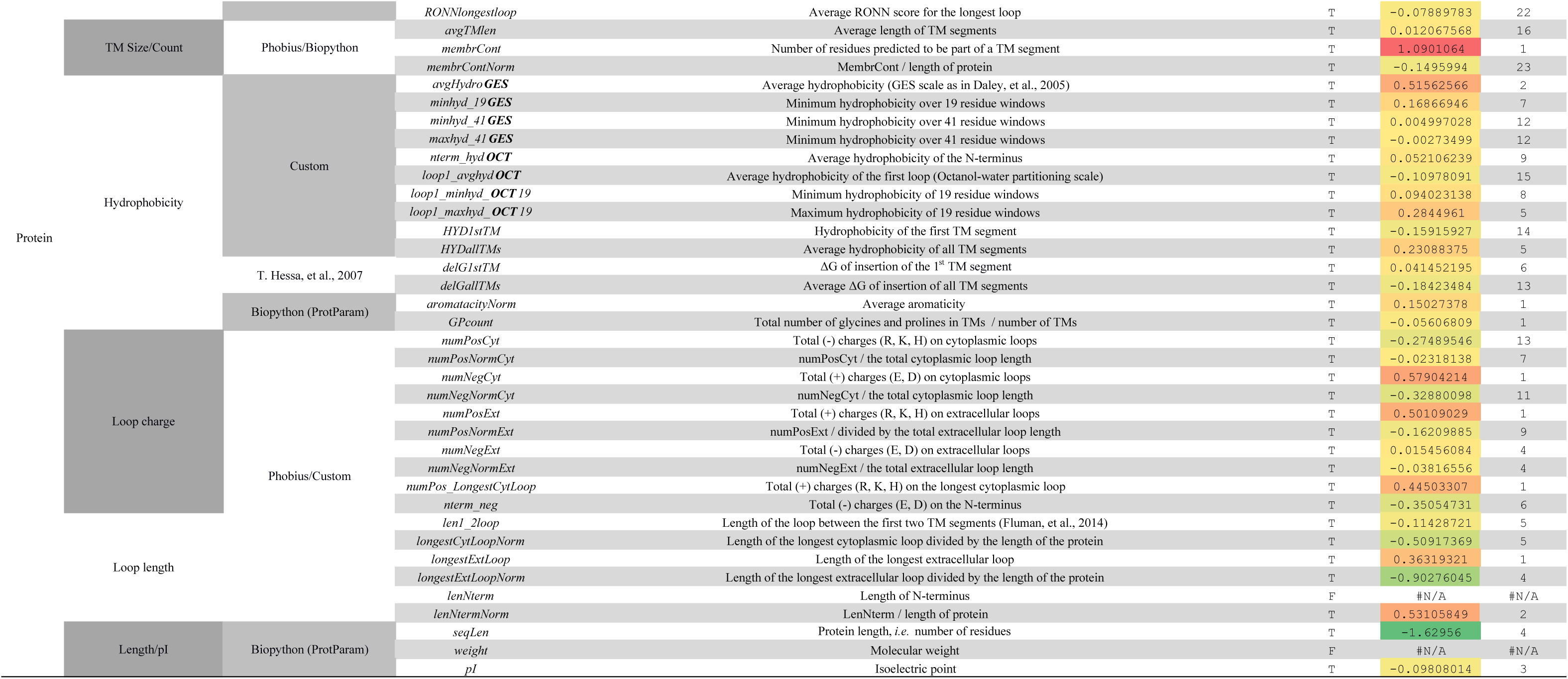
Sequence parameter weights and descriptions. Weights are presented after normalizing to the mean value for clarity. Features that were calculated but removed in pre-processing are noted (Methods 3).

**Supplementary Table 2.**
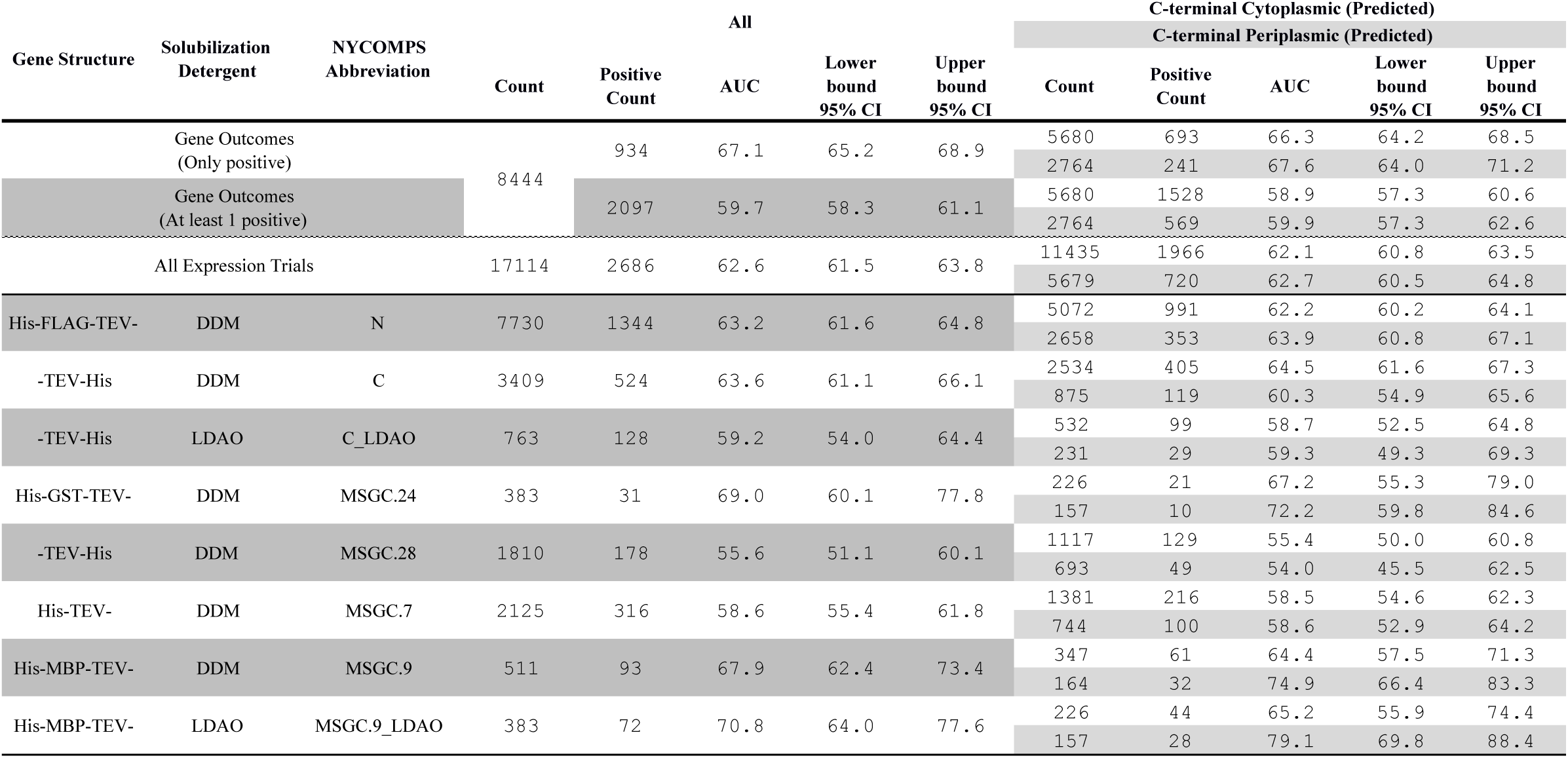
AUC values for the NYCOMPS dataset. AUC values and 95% confidence intervals are presented in summary, by expression condition, and by predicted C-terminal localization as well as for IMProve scores calculated without the most computationally expensive RNA secondary structure calculation.

**Supplementary Table 3. Predictive performances of the model across protein families.** The proteins and performances are with respect to those tested by NYCOMPS as summarized in Fig 2. This data is available in an interactive format at clemonslab.caltech.edu.

